# Antibiotic that inhibits *trans*-translation blocks binding of EF-Tu to tmRNA but not to tRNA

**DOI:** 10.1101/2023.06.09.544387

**Authors:** Neeraja Marathe, Ha An Nguyen, John N. Alumasa, Alexandra B. Kuzmishin Nagy, Michael Vazquez, Christine M. Dunham, Kenneth C. Keiler

## Abstract

*Trans-*Translation is conserved throughout bacteria and is essential in many species. High-throughput screening identified a tetrazole-based *trans*-translation inhibitor, KKL-55, that has broad-spectrum antibiotic activity. A biotinylated version of KKL-55 pulled down Elongation Factor Thermo-unstable (EF-Tu) from bacterial lysates. Purified EF-Tu bound KKL-55 *in vitro* with a *K_d_* = 2 µM, confirming a high-affinity interaction. An X-ray crystal structure showed KKL-55 binds in domain 3 of EF-Tu, and mutation of residues in the binding pocket abolished KKL-55 binding. RNA binding assays *in vitro* showed that KKL-55 inhibits binding between EF-Tu and tmRNA, but not between EF-Tu and tRNA. These data demonstrate a new mechanism for inhibition of EF-Tu function and suggest that this specific inhibition of EF-Tu•tmRNA binding is a viable target for antibiotic development.

**IMPORTANCE:** EF-Tu is a universally conserved translation factor that mediates productive interactions between tRNAs and the ribosome. In bacteria, EF-Tu also delivers tmRNA-SmpB to the ribosome during *trans*-translation. We report the first small molecule, KKL-55, that specifically inhibits EF-Tu activity in *trans*-translation without affecting its activity in normal translation. KKL-55 has broad-spectrum antibiotic activity, suggesting that compounds targeted to the tmRNA-binding interface of EF-Tu could be developed into new antibiotics to treat drug-resistant infections.

## INTRODUCTION

Antibiotic resistance is a critical public health concern due to the emergence and spread of multidrug-resistant infections. New therapeutic targets are urgently needed, and *trans*-translation is a promising pathway that is ubiquitous in bacteria but is not present in animals or humans (1–3). Bacteria use *trans*-translation as the primary ribosome rescue mechanism to recycle ribosomes that stall on nonstop mRNAs (truncated mRNAs that lack a stop codon) (4). *trans*-Translation is essential for survival in many pathogens such as *Mycobacterium tuberculosis, Shigella flexneri, Helicobacter pylori* and *Neisseria gonorrhoeae* (5–8), so targeting this pathway could be a promising strategy to combat a wide range of microbial threats classified as urgent and serious by the CDC. When ribosomes reach the 3′ end of an mRNA without encountering an in-frame stop codon, release factors cannot terminate translation. Because the frequency of stalling on nonstop mRNAs is high (an estimated 2-4% in *Escherichia coli*), bacteria cannot survive unless they can rescue the stalled ribosomes (4, 9). *trans*-Translation rescues nonstop ribosomes by providing a tRNA and mRNA mimic called transfer-messenger RNA (tmRNA) in complex with the small protein SmpB. This complex enters the ribosome and allows it to resume translation and terminate on a reading frame within tmRNA, thereby rescuing the stalled ribosome (4). The tmRNA reading frame encodes a degradation signal that is appended to the nascent peptide chain, prompting rapid proteolysis of the released protein (10–13).

Using a high-throughput screen of 663,000 candidate compounds, a small group of molecules were found to be effective *trans*-translation inhibitors (1). These molecules inhibited *trans*-translation both *in vitro* and *in vivo*, and exhibited broad-spectrum antibiotic activity against *S. flexneri*, *Bacillus anthracis*, and *Mycobacterium smegmatis*. KKL-55, a tetrazole-based compound, was one of the molecules that prevented C-terminal tagging and subsequent proteolysis of the nascent polypeptide. KKL-55 was later shown to be bactericidal to *B. anthracis* vegetative cells, germinants, and spores *in vitro* and after *ex vivo* infection of macrophages (14). Similarly, the minimum inhibitory concentration (MIC) of KKL-55 for *F. tularensis* was comparable to that of tetracycline (15). Notably, neither spontaneous nor UV-induced mutants of *E. coli, S. flexneri*, or *N. gonorrhoeae* that were resistant to KKL-55 could be recovered (1, 2). Here, we show that KKL-55 inhibits *trans*-translation by binding EF-Tu.

EF-Tu is an essential and universally conserved GTPase that is critical for protein synthesis. EF-Tu delivers aminoacyl-tRNAs (aa-tRNA) to the ribosomal aminoacyl (A) site during the elongation step of translation. Once the aa-tRNA docks to a cognate ribosomal A-site codon, EF-Tu exerts its GTPase activity, resulting in a conformational change that releases EF-Tu•GDP from the ribosome, and permits transfer of the nascent polypeptide from the P-site peptidyl-tRNA to the newly bound A-site aa-tRNA (16). Elongation factor thermo-stable (EF-Ts) binds EF-Tu and promotes exchange of GDP with GTP, regenerating the EF-Tu•GTP complex to deliver another aa-tRNA (17). EF-Tu contains three structural domains. Domain 1 (amino acids 1-200) contains the GTPase center. Domains 2 (amino acids 209 – 299) and 3 (amino acids 301 – 393) together bind aa-tRNA; domain 3 also interacts with EF-Ts to promote nucleotide exchange (18). During *trans*-translation, EF-Tu delivers alanyl-tmRNA-SmpB to the empty ribosomal A site. SmpB binds to the mRNA channel and the ribosomal decoding center and facilitates the accommodation of tmRNA so the nascent polypeptide chain can be transferred to its tRNA-like domain, and the tmRNA reading frame can be accepted as the new message (19).

Here, we perform structural and functional studies to understand how KKL-55 selectively inhibits *trans*-translation. We show that KKL-55 binds EF-Tu *in vitro*, and we solved a 2.2-Å X-ray crystal structure of *E. coli* EF-Tu co-crystallized with KKL-55, which shows KKL-55 binds to a highly conserved pocket in domain 3. We demonstrate that binding of KKL-55 has distinct effects on EF-Tu interaction with tRNA and tmRNA that explain the preferential inhibition of *trans*-translation.

## RESULTS

### EF-Tu is the molecular target of KKL-55

Previous modifications to the tetrazoyl benzamide KKL-55 suggested that large moieties could be added to the alkyl chain without impairing activity (14). To enable affinity purification of cellular molecules that bind to KKL-55, we designed KKL-201, an analog of KKL-55 that includes a biotin group (Fig. 1A). To ensure that KKL-201 retains the biochemical activity of KKL-55, and therefore is likely to bind the same target, we measured inhibition of *trans*-translation and translation by KKL-201 *in vitro*. *In vitro* transcription-translation assays using purified components from *E. coli* were programmed with a gene encoding DHFR with no stop codon at the 3′ end, and tmRNA-SmpB was added to the reactions. Transcription of the gene results in a nonstop mRNA, so translation and subsequent *trans*-translation results in a tagged DHFR protein (Fig. 1B). Like KKL-55, KKL-201 inhibited *trans*-translation in these reactions, resulting in a lower ratio of tagged to untagged protein. Neither KKL-55 nor KKL-201 inhibited protein synthesis when reactions were programmed with a gene encoding DHFR that included an in-frame stop codon (Fig. 1C). These results indicate that like KKL-55, KKL-201 specifically inhibits *trans*-translation and not normal translation.

**Figure 1.**
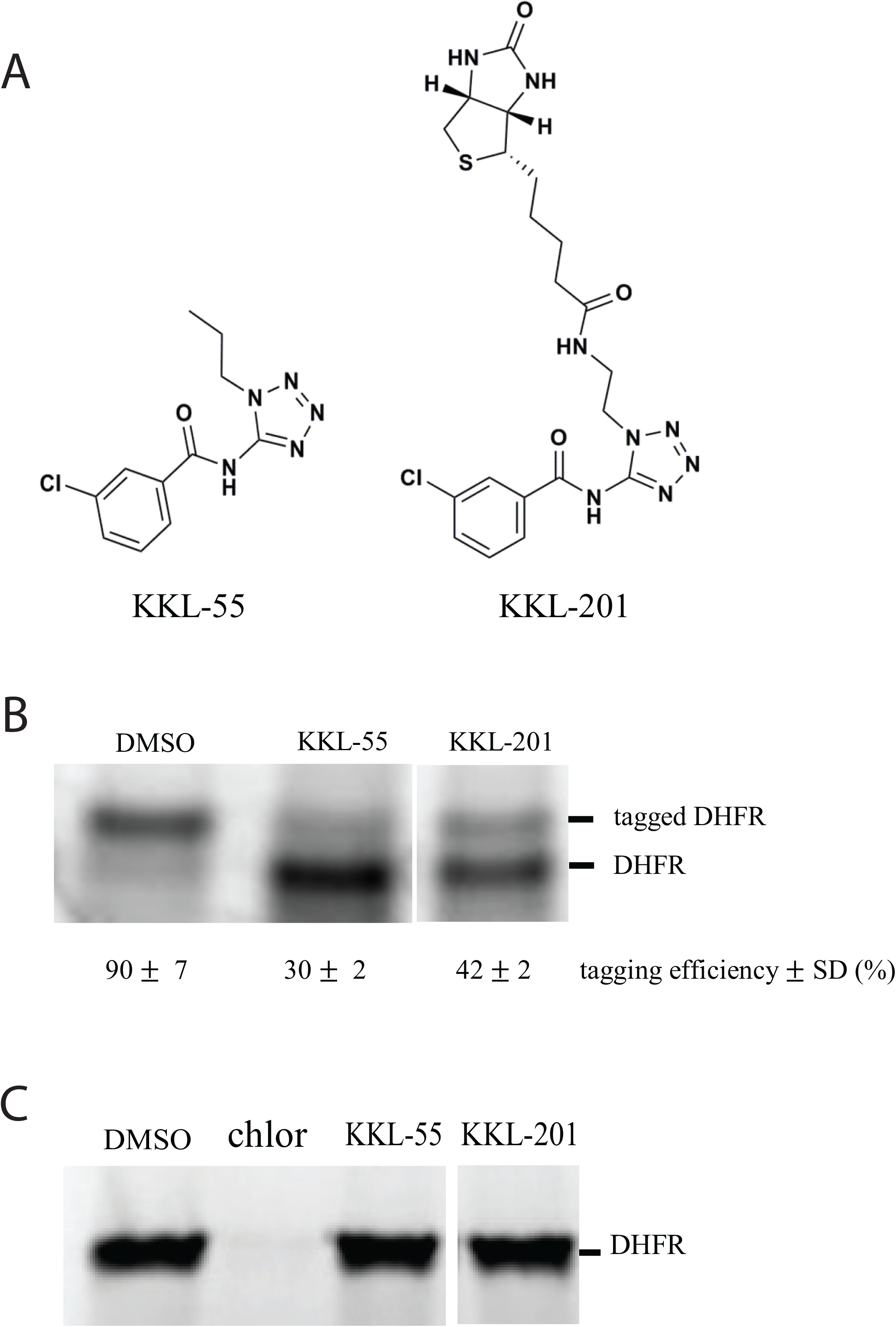
Tetrozoyl benzamides specifically inhibit *trans*-translation. A) Chemical structure of KKL-55 and KKL-201. B) *In vitro trans*-translation reactions in the presence of DMSO, 10 μM KKL-55, or 10 μM KKL-201. A gene encoding DHFR with no stop codon was transcribed and translated in the presence of tmRNA-SmpB and a tetrazoyl benzamide or vehicle. Protein products were labeled by incorporation of ^35^S-Met and analyzed by SDS-PAGE followed by autoradiography. A representative gel is shown with bands corresponding to DHFR and tagged DHFR indicated. The intensity of the DHFR and tagged DHFR bands were quantified and the tagging efficiency was calculated as the percentage of total DHFR protein in the tagged DHFR band. Mean tagging efficiency with standard deviation for at least 3 biological repeats is shown. C) *In vitro* translation reactions in the presence of DMSO, 100 μM chloramphenicol (chlor), 100 μM KKL-55, or 10 μM KKL-201. A gene encoding DHFR with stop codon was transcribed and translated as in (B) but without addition of tmRNA-SmpB. A representative gel is shown with the band corresponding to DHFR indicated.

To purify cellular molecules that bind to tetrazoyl benzamides, KKL-201 was incubated with a lysate from *B. anthracis* cells and the mixture was purified over Neutravidin resin. Affinity-purified proteins were visualized by SDS-PAGE, and inspection of the gel revealed 2 bands that appeared to be enriched in the eluate (Fig. 2A). Mass spectrometry identified these bands as pyruvate carboxylase and EF-Tu. Because pyruvate carboxylase contains a biotin co-factor which might allow it to bind to the Neutravidin resin independently of KKL-201, we investigated EF-Tu as a potential target for KKL-55.

**Figure 2.**
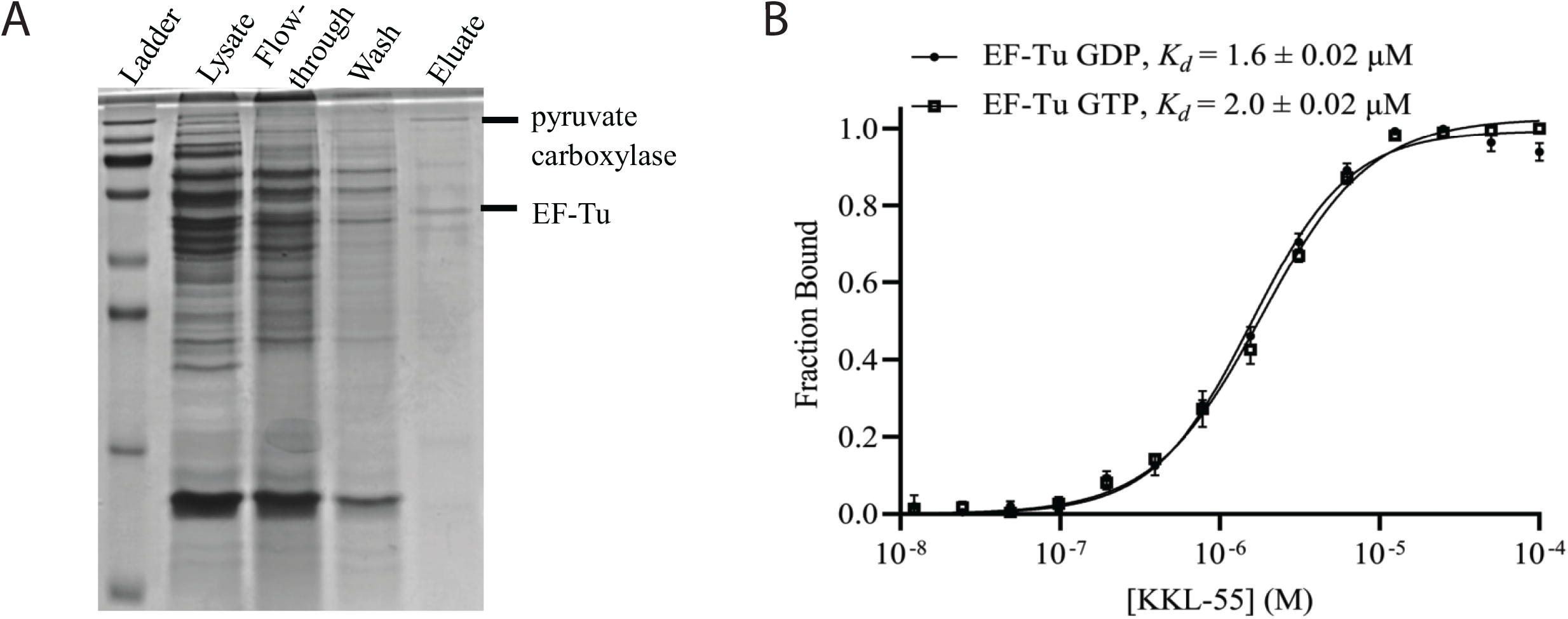
KKL-55 binds EF-Tu. A) SDS-PAGE analysis of KKL-201-affinity purified proteins from a *B. anthracis* lysate. Bands identified by mass spectrometry are indicated. B) MST assay for binding of EF-Tu and KKL-55 *in vitro*. Change in fluorescence was measured for fluorescently labelled EF-Tu•GTP or EF-Tu•GDP with different concentrations of KKL-55, and the fraction of EF-Tu bound to KKL-55 was calculated. Each point is the mean of 3 repeats with error bars indicating the standard deviation. The dissociation constant was calculated by non-linear curve fitting, and the mean dissociation constants with standard deviations for at least 3 repeats are shown.

### KKL-55 binds EF-Tu *in vitro*

We measured binding of KKL-55 with purified *E. coli* EF-Tu•GTP *in vitro* using microscale thermophoresis (MST) and observed concentration-dependent binding with an equilibrium binding constant (*K_d_*) of 2.0 µM (Fig. 2B). No binding was observed between EF-Tu and the structurally distinct *trans*-translation inhibitor KKL-35 (Fig. S2), which has been shown to bind to the ribosome (2, 8). EF-Tu•GDP bound KKL-55 with similar affinity as EF-Tu•GTP, indicating that the nucleotide state of EF-Tu is not important for KKL-55 binding (Fig. 2B). We measured the susceptibility of *E. coli ΔtolC* to KKL-55 using broth microdilution assays and the minimum inhibitory concentration (MIC) was 2.3 µM. The binding affinity for KKL-55 is therefore in the same range as the MIC, consistent with EF-Tu being the target responsible for growth inhibition by KKL-55.

### KKL-55 binds domain 3 of EF-Tu

To determine the binding site of KKL-55, we solved a 2.2 Å X-ray crystal structure of EF-Tu co-crystallized with KKL-55 (Figs. 3 & S4; Table 1). EF-Tu crystallized in the spacegroup P1 with two molecules per asymmetric unit. The structure was solved via molecular replacement using PDB code 6EZE as the starting model. The resulting difference F_o_-F_c_ density allows for the placement of KKL-55 into domain 3 of EF-Tu. Domains 2 and 3 adopt β-barrel structures and form the binding surface for aa-tRNAs (Fig. S3), and domain 3 is also the binding site for EF-Ts, critical for nucleotide exchange. In our structure, KKL-55 binds EF-Tu domain 3 in a pocket formed by residues Gly317, Arg318, His319, and Glu378 (Fig. 3B). Arg318 is at the bottom of the binding pocket and packs against KKL-55 from the tetrazole to the benzylchloride, with the positively charged guanidino group oriented toward the electronegative arene ring. Gly317 and His319 form one side of the binding pocket, and Glu378 forms electrostatic interactions with the tetrazole ring of KKL-55 on the other side of the pocket (Fig. 3C). These EF-Tu residues are broadly conserved (Fig. 3D), suggesting that KKL-55 is likely to bind EF-Tu from many bacterial species. The propyl group on KKL-55 extends out of the binding pocket, consistent with the observation that large substitutions at this position do not compromise activity (14). The arene ring of KKL-55 places the meta-chlorine towards the solvent (positioned away from the EF-Tu pocket), suggesting that substitutions could be made on this ring to improve activity.

**Figure 3.**
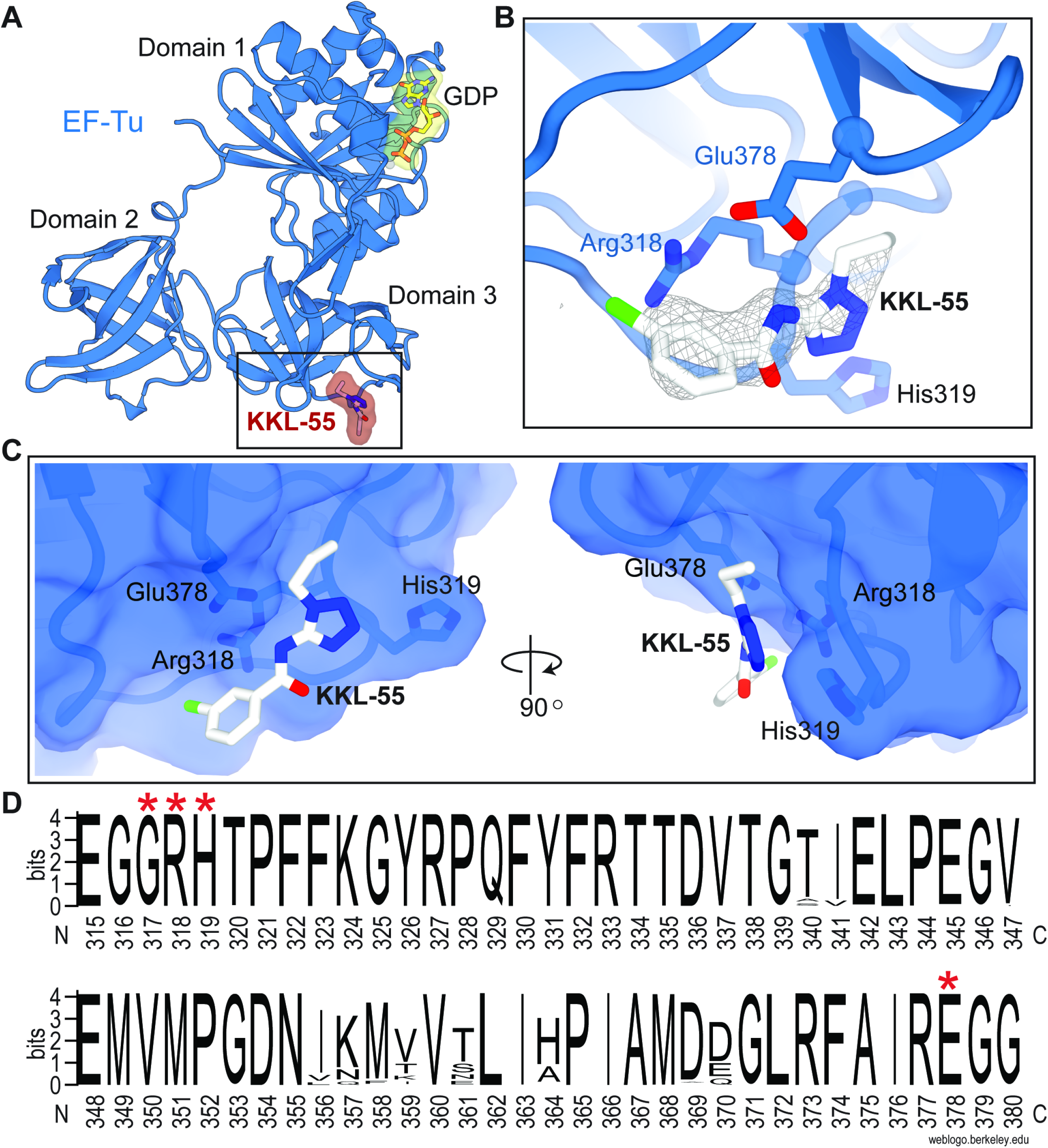
KKL-55 binds to a conserved region in domain 3 of EF-Tu. A) Overview of KKL-55 bound to Domain 3 of EF-Tu (blue) in the GDP-bound conformation. B) KKL-55 interacts with EF-Tu residues Glu378, Arg318 and His319. 2Fo-Fc electron density for KKL-55 is contoured at 1.0σ in gray mesh. C) Surface contour representation of EF-Tu and interacting residues Glu378, Arg318 and His319 surrounding KKL-55. D) Logo plot indicating the sequence conservation of KKL-55 residues 315-380 based on the analysis of 250 prokaryotic EF-Tu protein sequences homologous to *E. coli* K12 EF-Tu. Red asterisks indicate the strictly conserved residues of the KKL-55 binding pocket Gly317, Arg318, His319 and Glu378.

**Table 1:** Data collection and refinement statistics for the crystal structure of EF-Tu bound to KKL-55

|  |  |
| --- | --- |
| Data collection: |  |
| Space group | P1 |
| Resolution (Å) | 64.8 - 2.2 |
| Wavelength (Å) | 0.9792 |
| Cell dimensions: |  |
| a, b, c (Å) | 59.4, 65.2, 70.6 |
| $\alpha$, $\beta$, $\gamma$ (°) | 107.7, 98.8, 107.1 |
| Total reflections | 251,408 (22,029) |
| Unique reflections | 42,498 (3783) |
| $R_{merge}$ | 0.11 (0.88) |
| $R_{pim}$ | 0.05 (0.39) |
| $CC_{1/2}$ | 0.994 (0.777) |
| $CC^*$ | 0.998 (0.935) |
| $I/\sigma I$ | 9.79 (1.42) |
| Completeness (%) | 94.0 (84.3) |
| Redundancy | 5.9 (5.8) |
| Refinement: |  |
| Reflections used in refinement | 42,465 (3,783) |
| Reflections used for R-free | 2,038 (174) |
| $R_{work}/R_{free}$ | 0.25/0.27 |
| Ramachandran statistics |  |
| Favored (%) | 97.79 |
| Allowed (%) | 2.21 |
| Outliers (%) | 0 |
| RMS(bonds) | 0.008 |
| RMS(angles) | 1.25 |
| Clashscore | 2.58 |
| Average B-factor | 61.08 |
| macromolecules | 61.03 |
| ligands | 105.34 |
| solvent | 54.04 |

Based on the structure of EF-Tu-KKL-55, we made amino acid substitutions to Arg318, His319, and Glu378 and measured binding to KKL-55 using MST (Fig. 4). The R318A mutant had drastically lower affinity for KKL-55 (*K_d_* >30 µM), with a *K_d_* 15-fold higher than wild-type EF-Tu, consistent with the location of Arg318 in the KKL-55 binding pocket. Similarly, mutation of Arg318 to asparagine (R318N) dramatically reduced binding affinity (*K_d_* >20 µM). However, the R318K mutant showed only a modest decrease in affinity (*K_d_* = 2.6 µM), suggesting that the size and charge of the lysine residue allowing it to form similar interactions with KKL-55 as arginine at 318. The His319A had little impact on binding (Fig. 4), suggesting that the α-carbon of H319 is sufficient to form the pocket and the imidazole group is less important. Surprisingly, despite the electrostatic contacts between Glu378 and the tetrazole of KKL-55, the Glu378A mutation also had little effect on binding affinity (Fig. 4). The reason Glu378 appears to have no energetic contribution to binding is unclear, but in EF-Tu structures without KKL-55, Glu378 is oriented away from the KKL-55 binding pocket. Binding of KKL-55 results in reorientation of Glu378 and compensating movement of the peptide backbone of residues 379 and 380 away from the pocket. This structural change might cost approximately the same energy gained by the electrostatic interaction with KKL-55, resulting in no net binding energy. Simultaneous mutation of Arg318, His319, and Glu378 to alanine abolished detectable binding with KKL-55 (Fig. S5).

**Figure 4.**
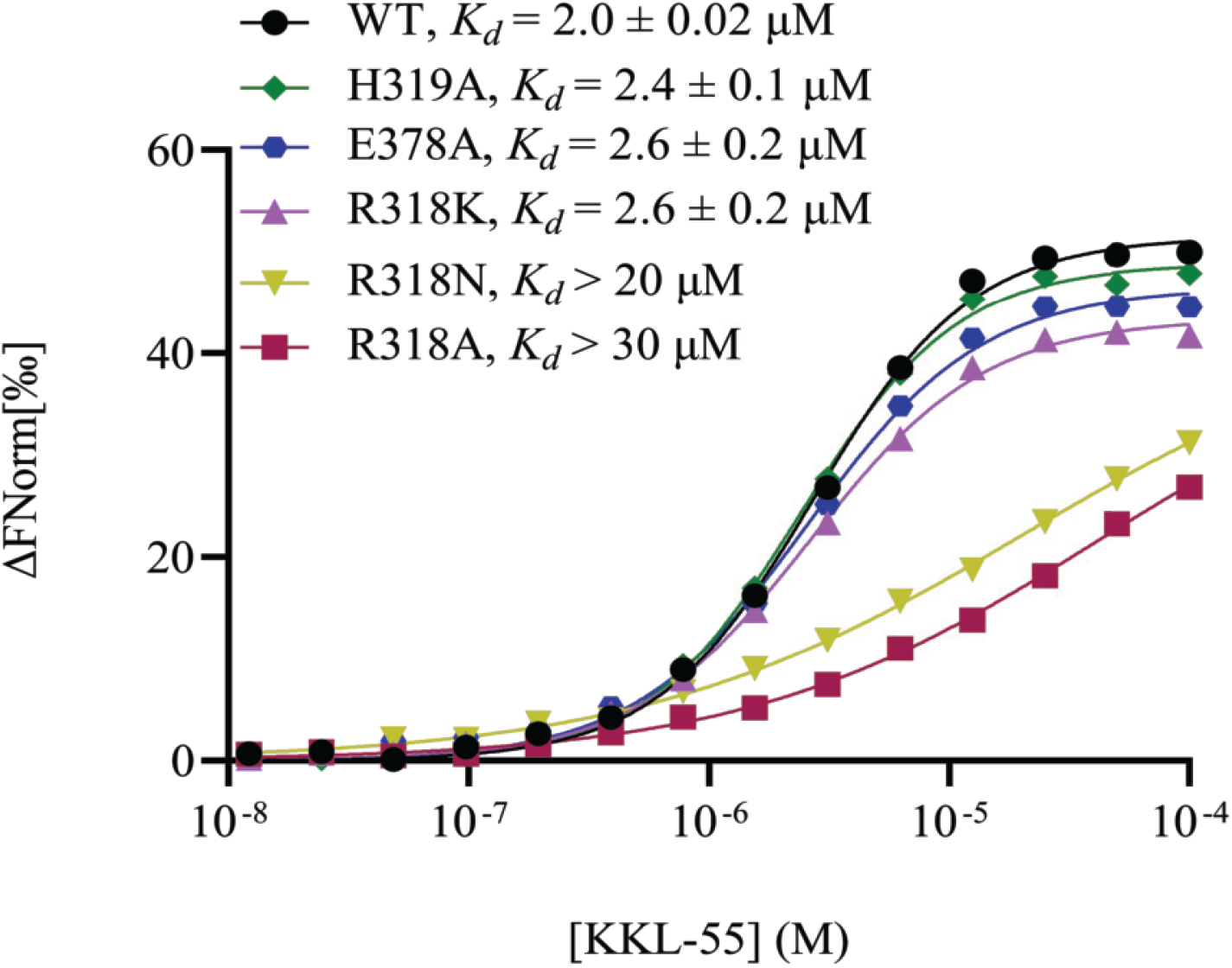
EF-Tu Arg318 is important for binding KKL-55. MST binding assays were performed with mutant versions of EF-Tu•GTP. One representative binding curve for each protein is shown. Data from three repeats were averaged and fit to sigmoidal function to determine the dissociation constant. The mean dissociation constant with standard deviation for at least 3 repeats is shown for each protein. Data for wild-type EF-Tu (WT) from Fig 2B is shown for comparison.

Because the R318A mutant has substantially lower binding with KKL-55, cells with this version of EF-Tu might be resistant to KKL-55. However, Arg318 is strictly conserved in EF-Tu proteins across bacteria (Fig. 3D), so it might be important for other functions of EF-Tu. We were not able to construct an *E. coli* strain that had EF-Tu R318A or R318N as the only copy of EF-Tu in the cell, and linked marker co-transduction experiments confirmed that cells were not viable when the only copy of EF-Tu was the R318A or R318N mutant (Fig. S6). Conversely, cells could survive with EF-Tu R318K as the only copy of EF-Tu (Fig. S6). Consistent with the tight binding of the R318K mutant and KKL-55 observed *in vitro*, cells with R318K had the same MIC for KKL-55 as wild type. These data indicate that Arg318 is important for viability in *E. coli*, and explain why mutations at this position do not confer resistance to KKL-55.

### KKL-55 binding alters EF-Tu binding to tmRNA

Because KKL-55 inhibits *trans*-translation but not normal translation, we hypothesized that KKL-55 alters or inhibits EF-Tu binding to tmRNA but has a smaller effect on the binding of EF-Tu to tRNA. To test this hypothesis, we used filter binding assays to measure the impact of KKL-55 on binding between EF-Tu•GTP and Ala-tmRNA or Ala-tRNA^Ala^. In the absence of KKL-55, EF-Tu•GTP bound to Ala-tmRNA with a *K_d_* = 0.75 µM and EF-Tu•GTP bound to Ala-tRNA^Ala^ with a *K_d_* = 0.28 µM (Fig. 5), similar to previously published data (20). In the presence of KKL-55, the binding affinity between EF-Tu•GTP and Ala-tmRNA was substantially reduced (Fig. 5A). Binding was weak enough that saturation could not be observed, but the *K_d_*> 10 µM. Conversely, KKL-55 had little effect on the binding affinity between EF-Tu•GTP and Ala-tRNA^Ala^ (Fig. 5B). This preferential inhibition explains how KKL-55 specifically inhibits *trans*-translation and not normal translation.

**Figure 5.**
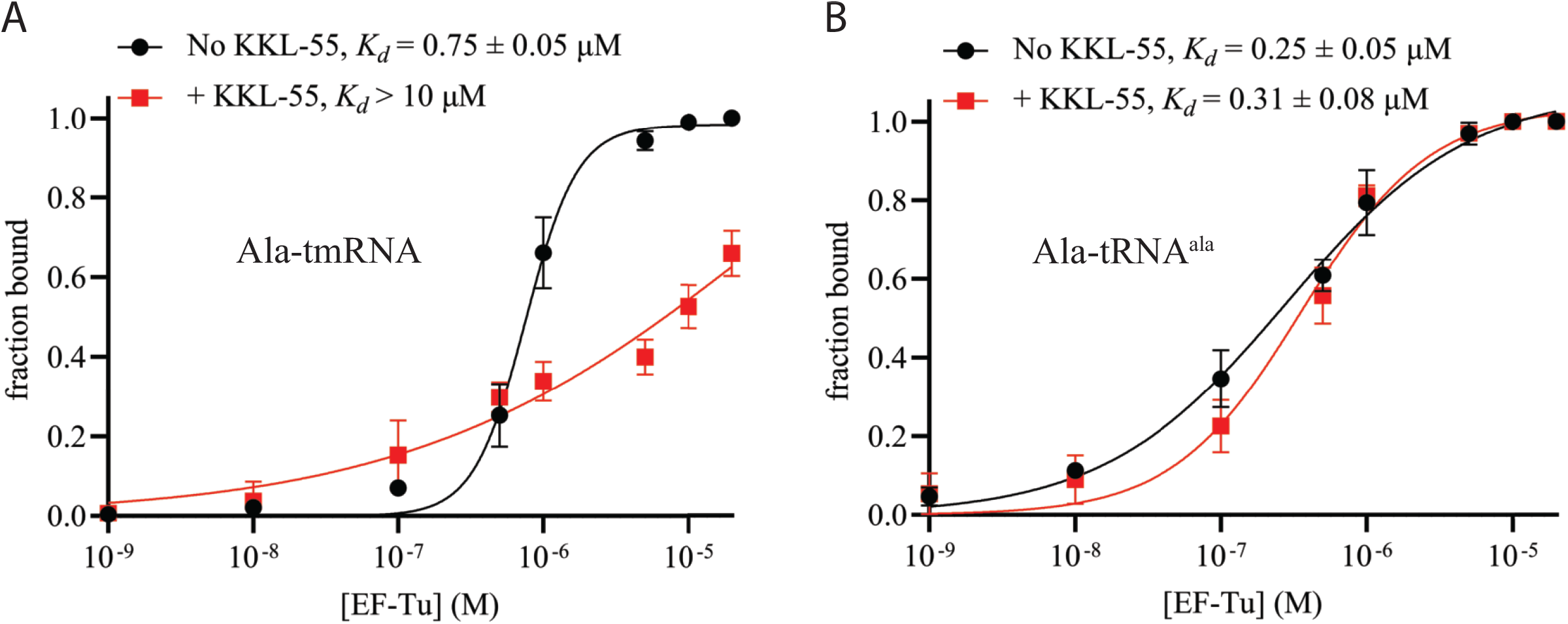
KKL-55 specifically inhibits EF-Tu binding with tmRNA. A) Filter binding assays were used to measure EF-Tu binding with Ala-tmRNA in the presence or absence of 25 μM KKL-55. Radiolabeled Ala-tmRNA was incubated with different concentrations of EF-Tu and bound Ala-tmRNA was separated using a nitrocellulose filter. The fraction of Ala-tmRNA bound was calculated and plotted versus EF-Tu concentration. Each point is the mean of 3 repeats with error bars indicating the standard deviation. The dissociation constant was calculated by non-linear curve fitting and the mean dissociation constant with standard deviation for at least 3 repeats is shown. B) Filter binding assays as in panel A to measure EF-Tu binding to Ala-tRNA^Ala^.

## DISCUSSION

The data shown here demonstrate that KKL-55 binds EF-Tu at a site distinct from other antibiotics. Binding of KKL-55 to this site dramatically decreases the affinity of EF-Tu for tmRNA. Because EF-Tu is required for tmRNA-SmpB to efficiently interact with the ribosome, inhibition of EF-Tu binding to tmRNA results in inhibition of *trans*-translation. The conservation of EF-Tu residues in the KKL-55 binding pocket suggests that KKL-55 should inhibit *trans*-translation in most bacterial species. This inhibition, together with the requirement for *trans*-translation in many bacteria, is consistent with the broad-spectrum antibiotic activity of KKL-55, although we have not excluded the possibility that KKL-55 also has other cellular targets that contribute to antibiotic activity.

Although the tRNA-like domain of tmRNA is very similar to a tRNA, EF-Tu binds to each RNA in subtly different ways at the elbow region, especially near the KKL-55 binding site in domain 3 (Fig. 6A). Overlaying EF-Tu•KKL-55 with tRNA and tmRNA reveals that KKL-55 binding to EF-Tu may alter interactions with each RNA substrate in slightly different ways. In the model of EF-Tu•tRNA•KKL-55, the arene end of KKL-55 would directly clash with the phosphate backbone of nucleotide 52 of the accepter arm of tRNA. Despite a predicted steric clash between KKL-55 and tRNA, KKL-55 has no effect on binding of EF-Tu to tRNA. The steric clash covers a single phosphate of the tRNA backbone, and presumably domain 3 of EF-Tu can move to prevent the clash without broadly disrupting other contacts with tRNA that make an important energetic contribution to binding. The same movement is clearly not possible when tmRNA is bound. Three factors may contribute to this difference. First, the total area of the steric clash is much larger for tmRNA (217.5 Å^2^ vs. 140.4 Å^2^; Fig. 6B). Second, the clash covers two nucleobases of tmRNA (nucleotides 339 and 340) near the middle of the tmRNA acceptor arm axis, which might require a much larger adjustment to avoid. Third, the paths of the tRNA and tmRNA backbones are different in this region, with tmRNA passing much closer to KKL-55 (Figs. 6A, S7). The closer proximity of tmRNA to EF-Tu and KKL-55 might sterically constrain movement of domain 3 or force disruption of more contacts between EF-Tu and tmRNA to avoid the clash with KKL-55. The tight binding of EF-Tu to tRNA in the presence of KKL-55 and the lack of inhibition of translation *in vitro* after addition of KKL-55 confirm that KKL-55 specifically disrupts *trans*-translation and not translation.

**Figure 6.**
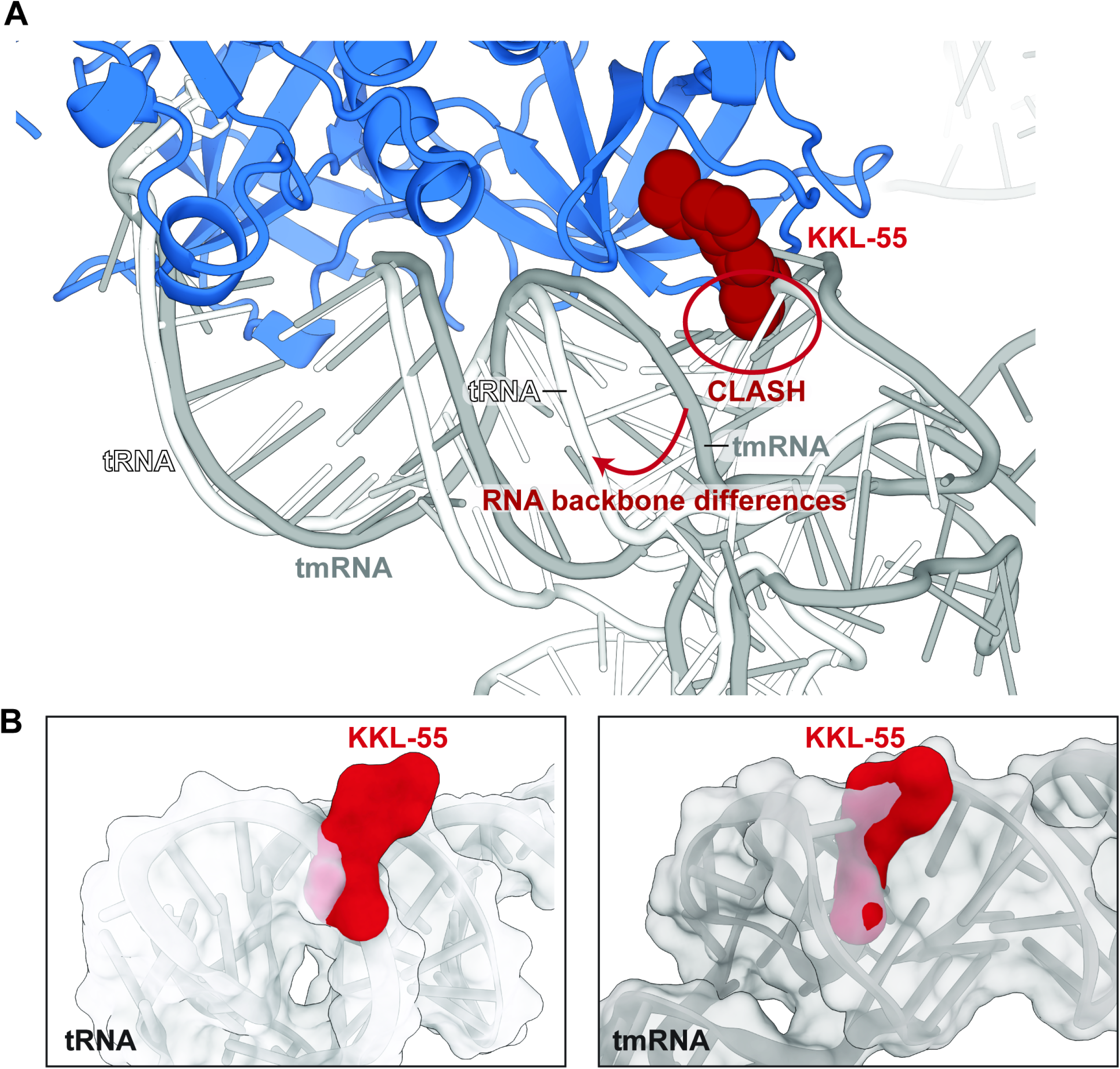
Comparison of EF-Tu binding interfaces with tRNA, tmRNA and KKL-55. A) Overlay of three structures of EF-Tu bound to tRNA (PDB code 1TTT), EF-Tu bound to tmRNA (PDB ID 7ABZ, EF-Tu in this structure not shown for clarity), and modeled KKL-55 as found in our structure in this study. Domains 2 and 3 of EF-Tu were used for alignments since Domain 1 is in the ‘open’, GDP-bound conformation. Differences between tRNA and tmRNA upon EF-Tu are localized to the acceptor arm and specifically tRNA nucleotides 59-63 and tmRNA nucleotides 348-351. The interaction between tRNA and EF-Tu at this region are further apart as compared to the tmRNA-EF-Tu interaction (as denoted by the red arrow and text). B) Surface representation of tRNA and tmRNA with KKL-55 (from PDBs above). The predicted clash area between is larger between KKL-55 and tmRNA (217.5 Å^2^) than KKL-55 and tRNA (140.4 Å^2^) consistent with stronger KKL-55 inhibition of tmRNA binding to EF-Tu.

The subtle but mechanistically significant differences in how KKL-55 affects EF-Tu interaction with tmRNA and tRNA parallels subtle mechanistic differences in how EF-Tu delivers tRNA and tmRNA to the ribosome. During normal translation, EF-Tu•GTP•aa-tRNA enters the A site and if there is no cognate base pairing between the tRNA anticodon and A-site codon, GTP hydrolysis is slow allowing EF-Tu•GDP•aa-tRNA to rapidly dissociate (21, 22). If the tRNA anticodon has cognate base pairing with the A-site codon, GTP hydrolysis on EF-Tu is rapid and EF-Tu•GDP dissociates, leaving the aa-tRNA accommodated in the A site (16). Delivery of aa-tmRNA-SmpB does not appear to be strictly coupled to GTP hydrolysis on EF-Tu. Experiments with translating ribosomes that have mRNA extending 3’ past the leading edge of the ribosome show they are not substrates for *trans*-translation (23). However, when EF-Tu•GTP•Ala-tmRNA-SmpB enters these non-substrate ribosomes, GTP hydrolysis is stimulated on EF-Tu even though Ala-tmRNA-SmpB is not accommodated into the A site (24). In addition, mutations in SmpB that reduce the rate of GTP hydrolysis on EF-Tu do not affect the affinity of Ala-tmRNA-SmpB for the ribosomal A site, suggesting that Ala-tmRNA-SmpB is released from EF-Tu more easily than tRNAs. Likewise, kirromycin appears to have differential activity between *trans-*translation and normal translation (25). Specifically, kirromycin does not inhibit EF-Tu•Ala-tmRNA-SmpB dissociation from the ribosome in the same way it inhibits EF-Tu•aa-tRNA dissociation during canonical elongation, supporting a different EF-Tu activation pathway for EF-Tu•Ala-tmRNA-SmpB as compared to EF-Tu•aa-tRNA (25). These data suggest that EF-Tu binds Ala-tmRNA and aa-tRNA differently during interaction with the ribosome, and the specific inhibition of EF-Tu•Ala-tmRNA binding by KKL-55 indicates that these differences are present before interaction with the ribosome as well.

EF-Tu is targeted by several antibiotics other than KKL-55. EF-Tu-binding antibiotics, also called elfamycins, primarily inhibit canonical translation and can be broadly categorized into two classes based on how they inhibit EF-Tu activity (26). The first group, consisting of pulvomycin and GE2270A, bind in the cleft between domains 1 and 2 of EF-Tu to prevent the essential conformation switch necessary for binding aa-tRNAs (27, 28). The second group, consisting of kirromycin and enacyloxin IIa, bind to the interface between domains 1 and 3 of EF-Tu, preventing EF-Tu from adopting its ‘open’ conformation that is necessary to dissociate from the ribosome (29, 30). The ability of KKL-55 to specifically target *trans*-translation by binding domain 3 of EF-Tu demonstrates a new mode of action by elfamycins and indicates that this binding site on EF-Tu could be targeted for development of new antibiotics to kill bacteria that require *trans*-translation.

## MATERIALS AND METHODS

### Reagents

Bacterial strains, plasmids, and primers used in this study are described in Tables S1-S3. Synthesis of KKL-201 is described in Scheme 1.

### *In vitro* translation and *trans*-translation assays

Translation assays were performed as previously described (1). Assays were set-up using PURExpress *in vitro* protein synthesis kit (New England Biolabs) with a cloned full-length DHFR template, and protein synthesis was monitored by incorporation of ^35^S-methionine. The inhibition of translation activity for each test compound was assessed with respect to the vehicle control from at least three independent assays. *In vitro trans*-translation was measured in a similar reaction that contained tmRNA-SmpB and employed a DHFR template missing two bases from the stop codon, as previously described (1). Due to the formation of a non-stop complex, tmRNA-SmpB can introduce an 11 amino acid tag on the DHFR protein which can be distinguished from the untagged DHFR protein on a SDS-PAGE gel. Test compounds were analyzed at 10 µM unless otherwise stated. Tagging efficiency was evaluated as the ratio of tagged DHFR to total DHFR from at least 3 repeats.

### Minimum Inhibitory Concentration (MIC) Assay

MIC assays were performed by broth microdilution according to Clinical and Laboratory Standards Institute guidelines for determining the antimicrobial activity of the compounds as described previously (1).

### Affinity Chromatography

*B. anthracis* cells were grown in 5 ml lysogeny broth (LB) at 37 °C overnight. This culture was grown in 1 L LB to a final OD_600_ ∼ 1.2. Cells were harvested by centrifugation at 14000 x *g* for 10 min, re-suspended in 25 ml lysis buffer (20 mM Tris pH 7.5, 2 mM β-mercaptoethanol (β-Me), 1 mg/ml lysozyme), lysed by sonication, and cell debris was removed by centrifugation at 28000 x *g*. The lysate was concentrated using a 10K Amicon ultra centrifugal filter. 500 µl concentrated lysate was added to 500 µl KKL-201 (400 µM) in binding buffer (100 mM Tris-HCl pH 8.0, 150 mM NaCl, 1 mM EDTA). This mixture was gently shaken at 4 °C for 2 h, added to NeutrAvidin agarose resin (ThermoFisher) equilibrated in binding buffer, incubated for 2 h, and loaded onto a column. The column was washed with 10 volumes binding buffer and bound proteins were eluted with 8 M guanidine hydrochloride. Fractions containing protein were combined, dialyzed against binding buffer, and concentrated using a 3K Amicon ultra centrifugal filter (Millipore). Purified proteins were separated on a 15% SDS-PAGE gel and visualized by staining with Coomassie Blue. Two enriched bands were excised and the proteins were identified by MS/MS mass spectrometry performed at the Protein & Nucleic Acid Facility (PAN) at Stanford University (Stanford, CA).

### Expression & Purification of *E. coli* EF-Tu

*E. coli* MG1655 pCA24N WT *tufA* (or mutant *tufA*) was grown in 1 L terrific broth (24 g/l yeast extract, 20 g/l tryptone, 4ml/L glycerol, 0.017 M KH_2_PO_4,_ 0.072 M K_2_HPO_4_) at 37 °C to OD_600_ = 0.6. 6His-tagged EF-Tu was overexpressed by growth in the presence of 1 mM isopropyl-thio-β-D-galactoside (IPTG) for 3 h. Cells at were harvested by centrifugation at 6953 x *g* for 10 min and stored at -80 °C. Harvested cells were resuspended in buffer (50 mM Tris-HCl pH 7.6, 60 mM NH_4_CI, 7 mM MgCl_2_, 7 mM β-Me, 15% (by vol.) glycerol, 10 µM guanosine diphosphate (GDP), 10 mM imidazole, 1 mM phenylmethylsµlfonyl fluoride), lysed by sonication, and debris was removed by centrifugation at 28000 x *g* for 15 min. The lysate was incubated with 750 µl HisPur Ni-NTA agarose resin (ThermoFischer) for 1 h, washed twice with buffer 1 (50 mM Tris-HCl pH 8.0, 60 mM NH_4_CI, 7 mM MgCl_2_, 7 mM β-Me, 15% (by vol.) glycerol, 10 µM GDP, 10 mM imidazole, 300 mM KCl), followed by two washes with buffer 2 (50 mM Tris-HCl pH 7.0, 60 mM NH_4_CI, 7 mM MgCl_2_, 7 mM β-Me, 15% (by vol.) glycerol, 10 µM GDP, 10 mM imidazole, 300 mM KCl), and loaded onto a column. Bound protein was eluted in buffer (50 mM Tris-HCl pH 7.6, 60 mM NH_4_CI, 7 mM MgCl_2_, 7 mM β-Me, 15% (by vol.) glycerol, 10 µM GDP, 500 mM imidazole), pooled and dialyzed in buffer (50 mM Tris-HCl pH 7.6, 60 mM NH_4_CI, 7 mM MgCl_2_, 7 mM β-Me, 15% (by vol.) glycerol, 10 µM GDP), and stored with 50% glycerol and 20 µM GDP at -20 °C.

### Apo-EF-Tu and EF-Tu•GTP preparation

Purified GDP bound 6His-tagged EF-Tu was incubated in buffer C (25 mM Tris-HCl pH 7.5, 50 mM NH_4_Cl, 10 mM EDTA) for 10 min at 37°C. Apo EF-Tu and GDP were separated by size exclusion chromatography on a Biorad ENRich SEC 70 column equilibrated in buffer F (25 mM Tris-HCl pH 7.5, 50 mM NH_4_Cl). GTP was added to 20 µM final concentration immediately after purification.

### EF-Tu purification for structural studies

EF-Tu was overexpressed from *E. coli* BL21-Gold (DE3) cells containing the pQE60-tufA-6xHis plasmid. Overnight cultures were grown in LB supplemented with 100 µg/ml ampicillin and 10 µg/ml tetracycline, diluted 1:500 into 4 L media. The bacterial culture was grown at 37 °C, 220 rpm to OD_600_ ∼0.6. Overexpression of EF-Tu was induced by the addition of 0.1 mM IPTG for 4 h at 37 °C. Cell pellets were harvested by centrifugation and lysed using an Emulsiflex C5 (Avestin) in lysis buffer (50 mM HEPES-KOH pH 7.6, 1 M NH_4_Cl, 10 mM MgCl_2_, 0.3 mg/ml lysozyme, 0.1% Triton X-100, 0.2 mM PMSF, 7 mM β-Me) for three passes. The cell lysate was clarified by centrifugation (30,392 x *g* at 4 °C for 90 min), then loaded on to a 5 ml Ni^2+^ HP HisTrap column (Cytiva) for affinity purification. EF-Tu was eluted using a 10-400 mM imidazole gradient (50 mM HEPES-KOH pH 7.6, 1 M NH_4_Cl, 10 mM MgCl_2_, 7 mM β-Me) and then loaded on a Superdex 200 HiLoad 16/60 Prep Grade column (Cytiva) in buffer (50 mM HEPES-KOH pH 7.6, 100 mM KCl, 10 mM MgCl_2_, 30% glycerol, 7 mM β-Me). The protein was dialyzed into the final storage buffer (30 mM Tris pH 8.0, 200 mM (NH_4_)_2_SO_4_, 1 mM MgCl_2_, 1 mM DTT, 10% glycerol) and then concentrated using a 10K Amicon Ultra centrifugal filter (Millipore). The protein concentration was determined by a Bradford assay and stored as aliquots at -80 °C.

### Structural determination of the EF-Tu•KKL-55 complex

Crystallization trials of 11 mg/ml EF-Tu were performed using the Phoenix protein crystallization robot (Arts Robbins Instruments) using a 400 nl drop size with a 1:1 ratio of protein to reservoir condition in Intelli-Plate 96-3 LVR plates. EF-Tu crystals grew in 30-35% polyethylene glycol monomethyl ether 5,000 (PEG 5K MME), 0.2 M (NH_4_)_2_(SO_4_), and 0.1 M MES pH 6.5. KKL-55 was dissolved in 100% DMSO and stored in the dark at -20 °C. The EF-Tu•KKL-55 complex was prepared by incubating 11 mg/ml (or 254 µM) EF-Tu with 0.2 mM KKL-55 for 30 min at room temperature. The mixture was then centrifuged to pellet any precipitation for 3 min at 20,627 x *g*, and the supernatant was used to set up sitting drops using 3.6 µl drop size in a 1:1 protein to reservoir ratio over 400 µl of reservoir volume at 20 °C. Needle-like crystals grew within 1-2 days and were flash frozen after being cryoprotected stepwise with solutions of 5/15/25% PEG 400, 0.2 M (NH_4_)_2_(SO_4_), 20% PEG 5K MME and 0.1 M MES pH 6.5. Datasets were integrated and scaled using XDS (31) and the Northeastern Collaborative Access Team (NE-CAT) RAPD system. The structure was phased using molecular replacement with the EF-Tu model from PDB code 6EZE (32), and the model was iteratively refined in PHENIX (33) and built in Coot (34). The KKL-55 model was generated in ChemDraw and refinement restraints generated using eLBOW in PHENIX (35). Feature enhanced maps were generated in PHENIX (36). Figures were generated in PyMOL. Surface area calculations were performed in ChimeraX (37).

### Conservation analysis of EF-Tu and KKL-55 binding pocket

Conservation analyses of the KKL-55 binding pocket were performed on ∼250 prokaryotic EF-Tu protein sequences identified using BLASTp with the sequence from 6EZE against the UniProtKb protein sequence database (38). Amino acid sequences were downloaded from UniProt and duplicate sequences and those of unknown function were removed. Protein sequences were aligned using MUSCLE (39). A frequency logo was created from aligned EF-Tu sequences using WebLogo (40). ConSurf analysis was performed with conservation score determined by Bayesian inference (41) (Fig. S8).

### Site-directed mutagenesis of EF-Tu

All mutants were constructed using HiFi assembly (New England Biolabs). Two PCR products were generated for each mutant using primer pairs listed in Table S3, with pCA24N-His6-tufA as template. The PCR products were assembled with the pCA24N-His6-tufA that had been digested with BamHI and NotI.

### Microscale thermophoresis binding assays

300 nM 6His-tagged EF-Tu was incubated with 75 nM Red Tris NTA dye Generation 2 (NanoTemper Technologies) in binding buffer (287 mM NaCl, 2.7 mM KCl, 10 mM Na_2_HPO_4_, 1.8 mM KH_2_PO_4_, 0.01 % Tween 20) for 30 min followed by centrifugation at 21,000 x *g* for 10 min. The EF-Tu and dye complex was added to KKL-55 in a 1:1 ratio and incubated at room temperature for 2 h, and MST was measured in a Monolith NT.115 (NanoTemper Technologies). Plots of change in fluorescence vs. the concentration of KKL-55 were fit to the hyperbolic function y = c/(1 + *Kd*/x) to obtain the apparent binding constant.

### Phage Transduction

Tet^R^-*tufA::Kan* linked marker strain (KCK575) was constructed via recombination (42) of Tet^R^ into the chromosome of *tufA* deletion strain, 15.6 kb away from *tufA::Kan*, and a P1 phage lysate (43) was prepared and used to transduce Δ*tolC* Δ*tufB* cells harboring pCA24N-His6-tufA, pCA24N-His6R318A or pCA24NHis6R318N. Cells were plated on LB with oxytetracycline, chloramphenicol and IPTG to select for transductants. The resulting colonies were tested for kanamycin resistance to determine the co-transduction frequency.

### Filter Binding Assays

[α-^32^P]-ATP body labelled tRNA^Ala^ and tmRNA was prepared by *in vitro* transcription (44) and aminoacylated using purified AlaRS as previously described (45). 1 ml master binding mix was prepared with 0.1 nM [α-^32^P]-Ala-tRNA^Ala^ or [α-^32^P]-Ala-tmRNA in binding buffer (44 mM KH_2_PO_4_, 6 mM K_2_HPO_4_, 10 mM MgCl_2,_ 50 µg/ml BSA pH 5.5). 16 µl of each concentration of 6His-tagged EF-Tu was mixed with 34 µl RNA binding mix and incubated for 15 s at room temperature. The reaction mix was passed under vacuum through 0.45 µm nitrocellulose membrane filter (Millipore) and positively charged nylon membrane (Amersham) presoaked with wash buffer (44 mM KH_2_PO_4_, 6 mM K_2_HPO_4_, 10 mM MgCl_2_ pH 5.5). The membranes were washed, dried, and radioactivity was determined by scintillation counting. Where appropriate, 25 µM KKL-55 was pre-incubated with 6His-tagged EF-Tu. Plots of fraction EF-Tu bound vs. concentration of EF-Tu were fit to the hyperbolic function y = c/(1 + *Kd*/x) to obtain the apparent equilibrium binding constants.

## ACKNOWLEDGEMENTS

We thank Keiler and Dunham laboratory members for their critical reading of the manuscript, and Dongxue Wang for technical expertise. Support for this work was provided by NIH IRACDA 2K12GM000680 (ABKN) and NIH R01GM121650 (KCK, CMD). This work is based upon research conducted at the Northeastern Collaborative Access Team beamlines, which are funded by the National Institute of General Medical Sciences from the National Institutes of Health (P30 GM124165). This research used resources of the Advanced Photon Source, a U.S. Department of Energy (DOE) Office of Science User Facility operated for the DOE Office of Science by Argonne National Laboratory under Contract No. DE-AC02-06CH11357.

---

**Scheme 1.**
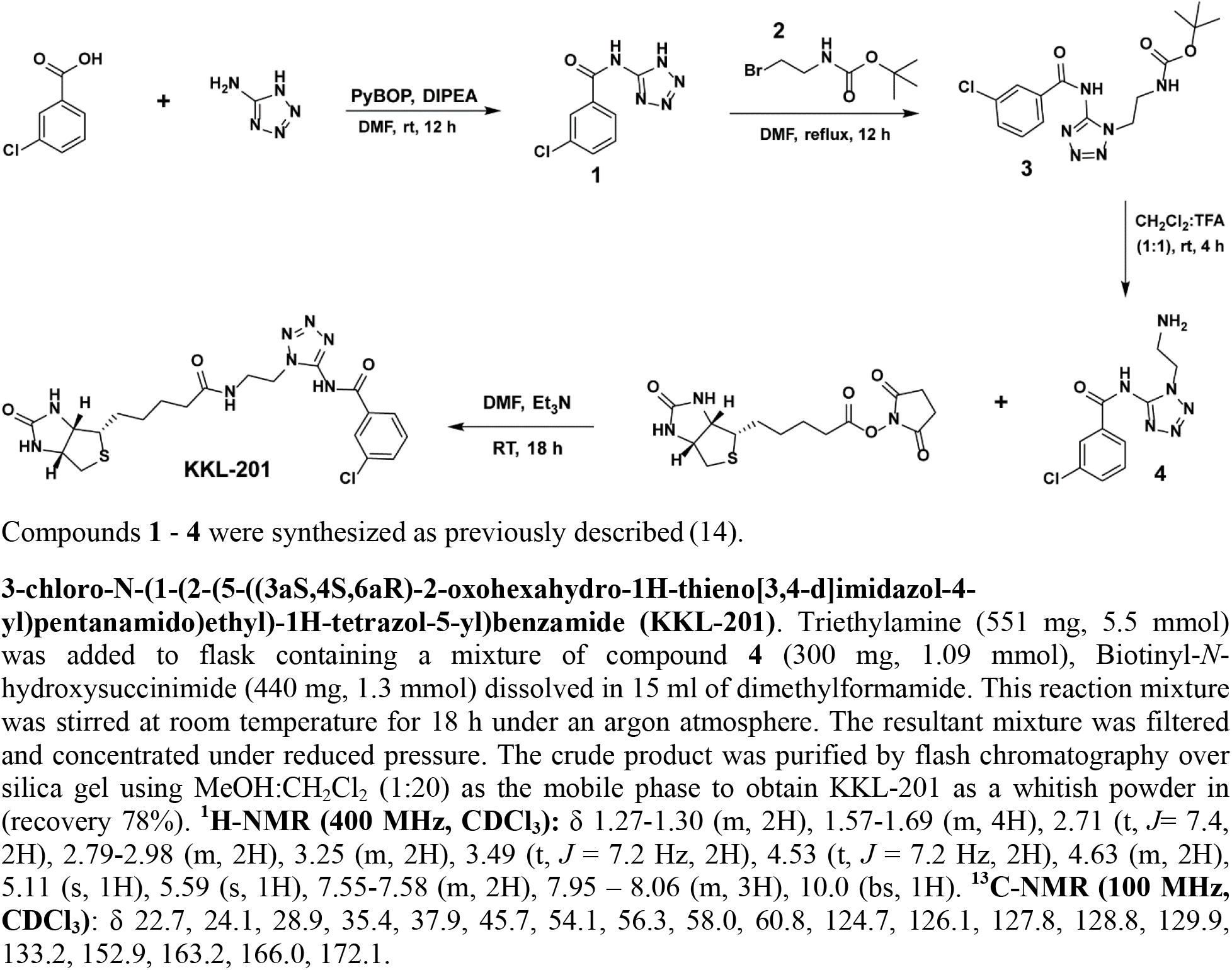
Synthesis of 3-chloro-N-(1-(2-(5-((3aS,4S,6aR)-2-oxohexahydro-1H-thieno[3,4-d]imidazol-4-yl)pentanamido)ethyl)-1H-tetrazol-5-yl)benzamide (KKL-201) General: All organic solvents and reagents were purchased from Sigma-Aldrich (St. Louis, MA). unless otherwise stated. Chloroform-*d* was purchased from Cambridge Isotope Laboratories (Andover, MA). Nuclear Magnetic Resonance analyses were conducted on a 400 MHz Bruker spectrophotometer.

## FIGURE LEGENDS

**Figure S1. EF-Tu is not enriched in a mock affinity purification.** Affinity purification was performed as described in Figure 2A but without the addition of KKL-201.

**Figure S2. EF-Tu does not bind to structurally distinct trans-translation inhibitor.** A) Chemical structure of KKL-35. B) MST assay for binding of EF-Tu and KKL-35 *in vitro*. Change in fluorescence was measured for fluorescently labelled EF-Tu•GTP with different concentrations of KKL-35. The dissociate constant was calculated by non-linear curve fitting, and the mean dissociation constants with standard deviations for at least 3 repeats are shown.

**Figure S3. Domains 2 and 3 of EF-Tu (blue) binds tRNA (white) directly.** Figure was generated from PDB code 1TTT.

**Figure S4. Structural determination of EF-Tu binding to KKL-55.** A) Needle-like crystals ready for data collection grew in 1-2 days. B) Photo of the crystal from which this dataset was collected from. C) Cartoon representation of the asymmetric unit of the crystal containing 2 EF-Tu molecules. The overlay of gray mesh represents the 2F_o_-F_c_ electron density map contoured at 1.0σ.

**Figure S5. EF-Tu Arg318 is important for EF-Tu•GDP binding KKL-55.** MST binding assays were performed with mutant versions of EF-Tu•GDP. One representative binding curve for each protein is shown. Each point is the mean of 3 repeats with error bars indicating the standard deviation. Data from three repeats were averaged and fit to sigmoidal function to determine the dissociation constant. The mean dissociation constant with standard deviation for at least 3 repeats is shown for each protein. Data for wild-type EF-Tu (WT) from Fig 2B is shown for comparison.

**Figure S6. R318A and R318N mutants do not support viability of *E. coli*.** A) Cartoon depicting the co-transduction experiment. P1 transduction was used to try to delete both chromosomal genes expressing EF-Tu from the cell and express wild-type or mutant EF-Tu from a plasmid. B) Table with co-transduction frequency results. C) Representative plates for results from panel B. Tet plates (left) and Kan plates (right) for R318A (top) and R318N (bottom) mutants. One positive control (*ΔtufB* pWT*tufA*) was used on one Kan plate (yellow box, top right plate).

**Figure S7. Alignment of a 70S pre-accommodation EF-Tu-tRNA structure with the EF-Tu-KKL-55 structure.** A) A 70S pre-accommodation EF-Tu structure containing an A*/T tRNA state (PDB code 6WD2) aligned with the EF-Tu-KKL-55 structure reveals similar interactions with the modeled KKL-55. Comparison of the structures reveals that domain 1 undergoes a large rotation upon tRNA binding while domains 2 and 3 are very similar. The EF-Tu conformation in the pre-accommodation state is similar to both an EF-Tu-tRNA structure off the ribosome (PDB code 1TTT) and a 70S-preaccommodated EF-Tu-tmRNA structure (PDB code 7ABZ; these structures are not shown for clarity). B) Comparison of only domains 2 and 3 emphasize the substantial similarity between structures.

**Figure S8: Logo plot of EF-Tu.** Logo plot as in Fig. 3D showing the entire EF-Tu protein sequence.

---

**Table S1:** Strains used in this study

| Strain | Description | Source or Reference |
| --- | --- | --- |
| *B. anthracis* Sterne | pXO1<sup>+</sup>, pXO2<sup>-</sup> | Gift from Turnbough Lab |
| *E. coli* DH5α | cloning strain | New England Biolabs |
| JW5503 | *tolC::Kan* | (46) |
| *E. coli* Δ*tolC*Δ*tufB* | Markerless with *tolC* and *tufB* deletion | This study |
| KCK571 | *E. coli* Δ*tolC*Δ*tufB* Δ*tufA::Kan* with WT *tufA* on pCA24N | This study |
| KCK572 | *E. coli* Δ*tolC*Δ*tufB* Δ*tufA::Kan* with pCA24NR318K | This study |
| KCK573 | *E. coli* Δ*tolC*Δ*tufB* Δ*tufA::Kan* with pCA24NH319A | This study |
| KCK574 | *E. coli* Δ*tolC*Δ*tufB* Δ*tufA::Kan* with pCA24NE378A | This study |
| JW3301 | *tufA::Kan* | (46) |
| KCK559 | DH5α with pCA24NR318A | This study |
| KCK560 | DH5α with pCA24NR318K | This study |
| KCK561 | DH5α with pCA24NR318N | This study |
| KCK562 | DH5α with pCA24NH319A | This study |
| KCK563 | DH5α with pCA24NE378A | This study |
| KCK564 | DH5α with pCA24NR318AH319AE378A | This study |
| KCK565 | *E. coli* Δ*tolC*Δ*tufB* with WT *tufA* on pCA24N | This study |
| KCK566 | *E. coli* Δ*tolC*Δ*tufB* with pCA24NR318A | This study |
| KCK567 | *E. coli* $\Delta tolC\Delta tufB$ with pCA24NR318K | This study |
| KCK568 | *E. coli* $\Delta tolC\Delta tufB$ with pCA24NR318N | This study |
| KCK569 | *E. coli* $\Delta tolC\Delta tufB$ with pCA24NH319A | This study |
| KCK570 | *E. coli* $\Delta tolC\Delta tufB$ with pCA24NE378A | This study |
| JW3301 | tufA-6XHis | (47) |
| KCK575 | Tetracycline resistance marker linked to *tufA*::*Kan* in genome | This study |

**Table S2:**
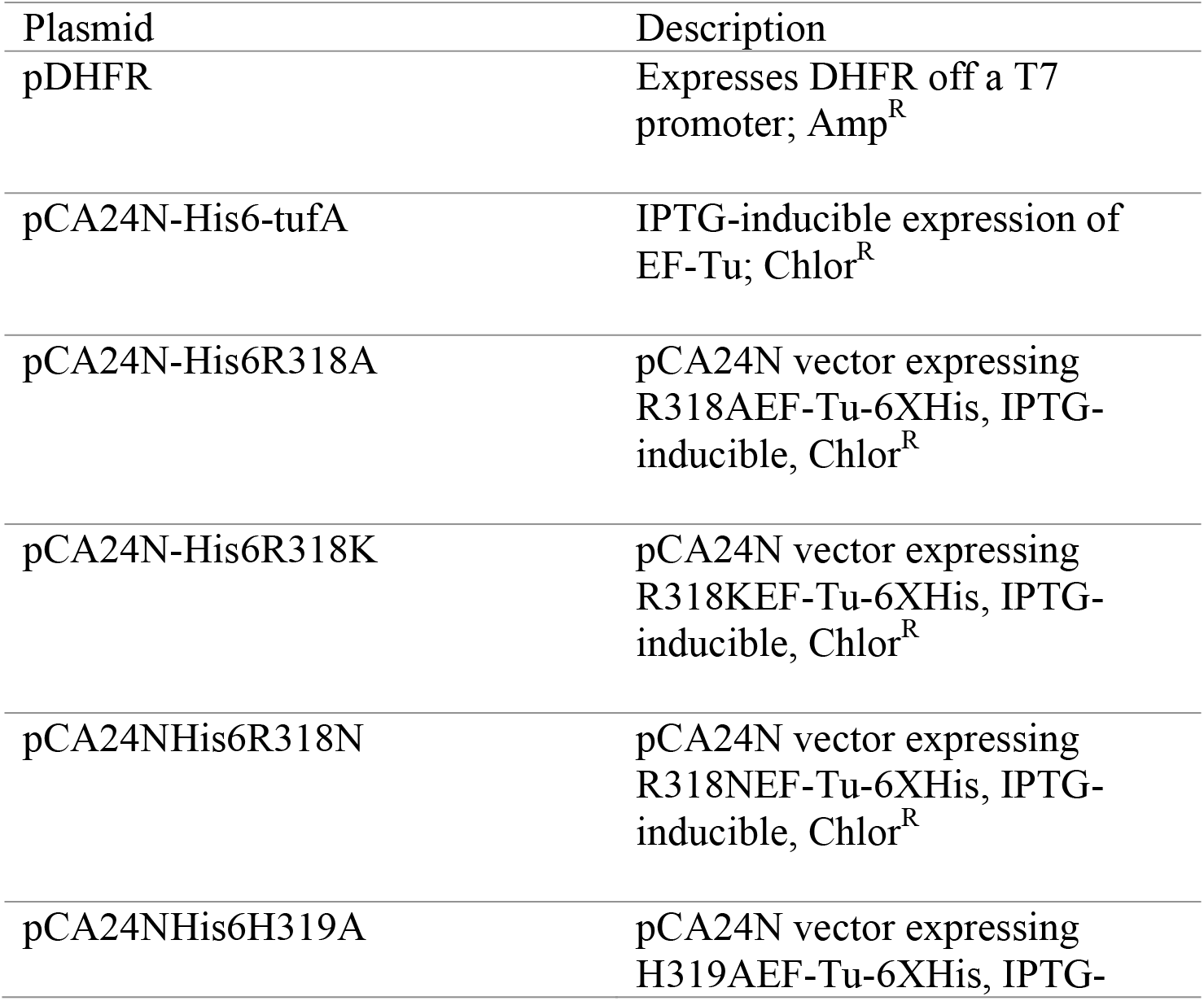
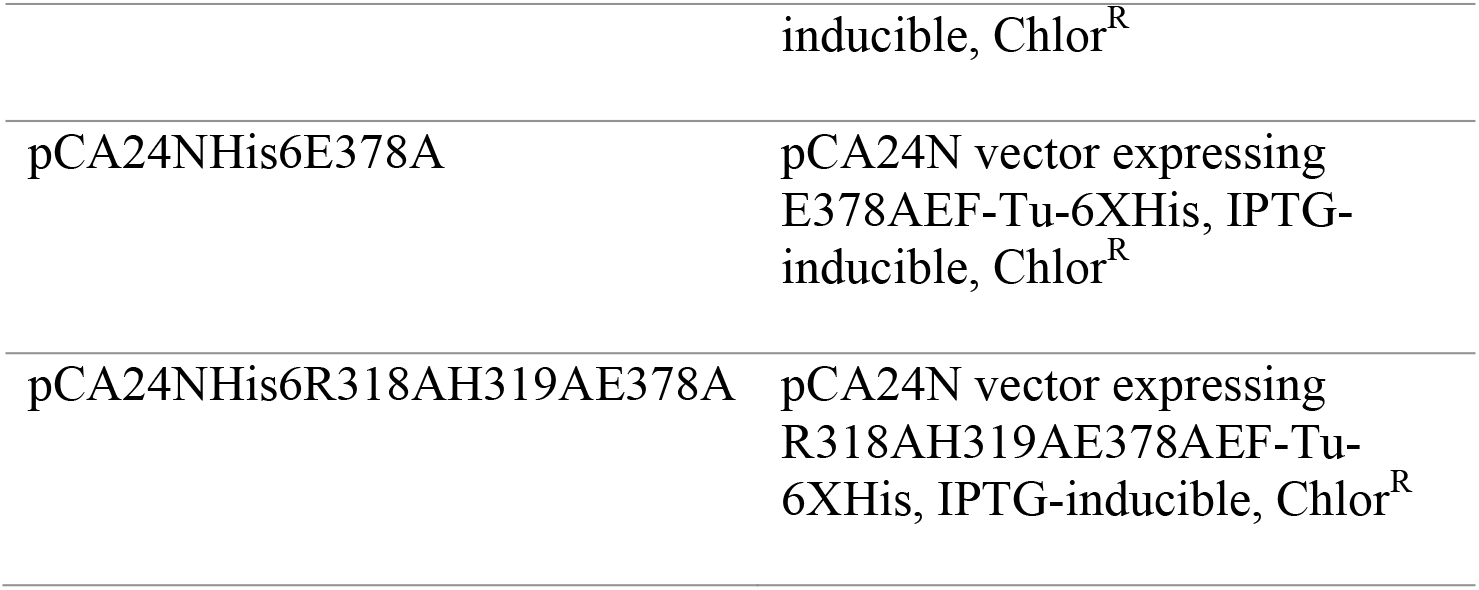
Plasmids used in this study

**Table S3:**
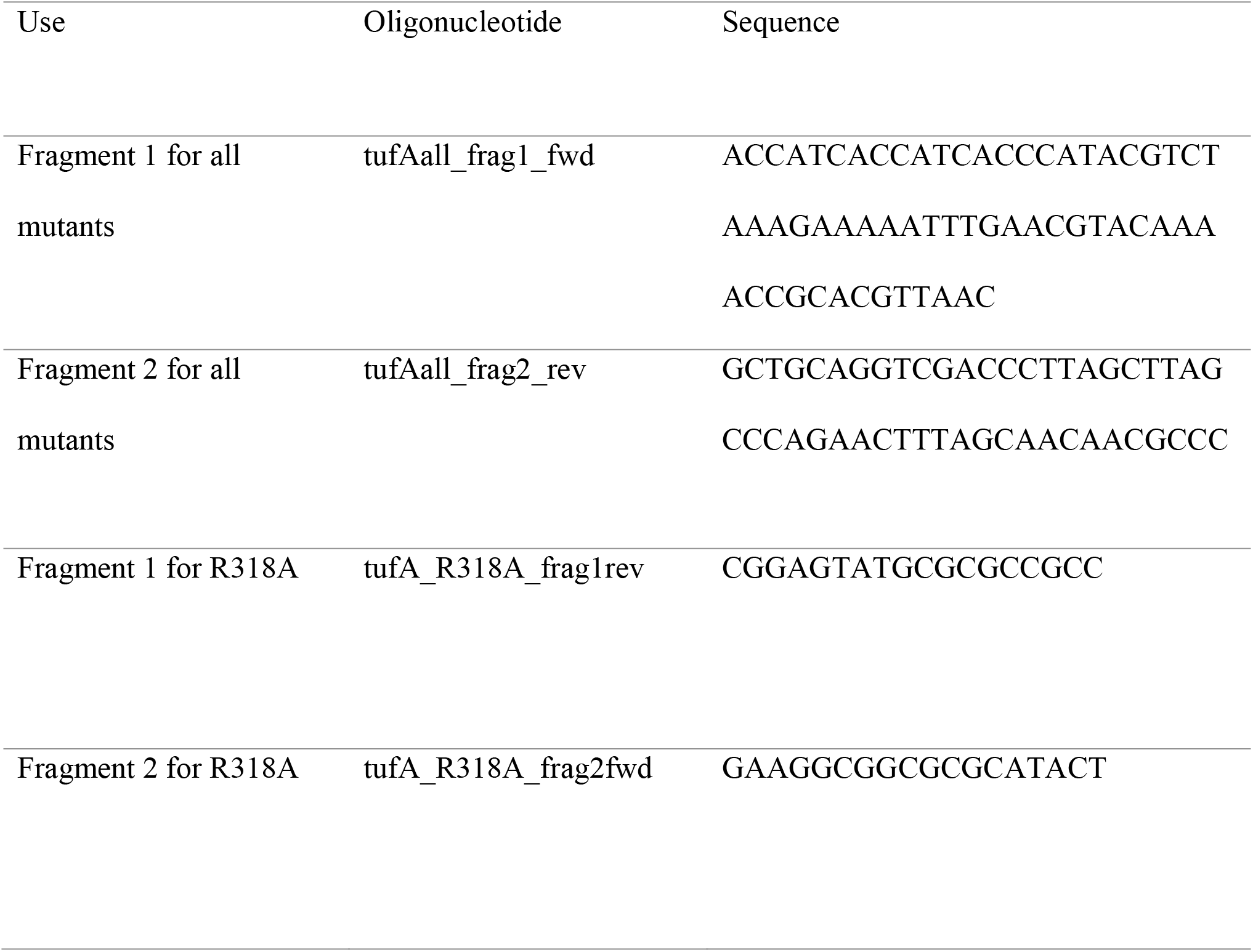
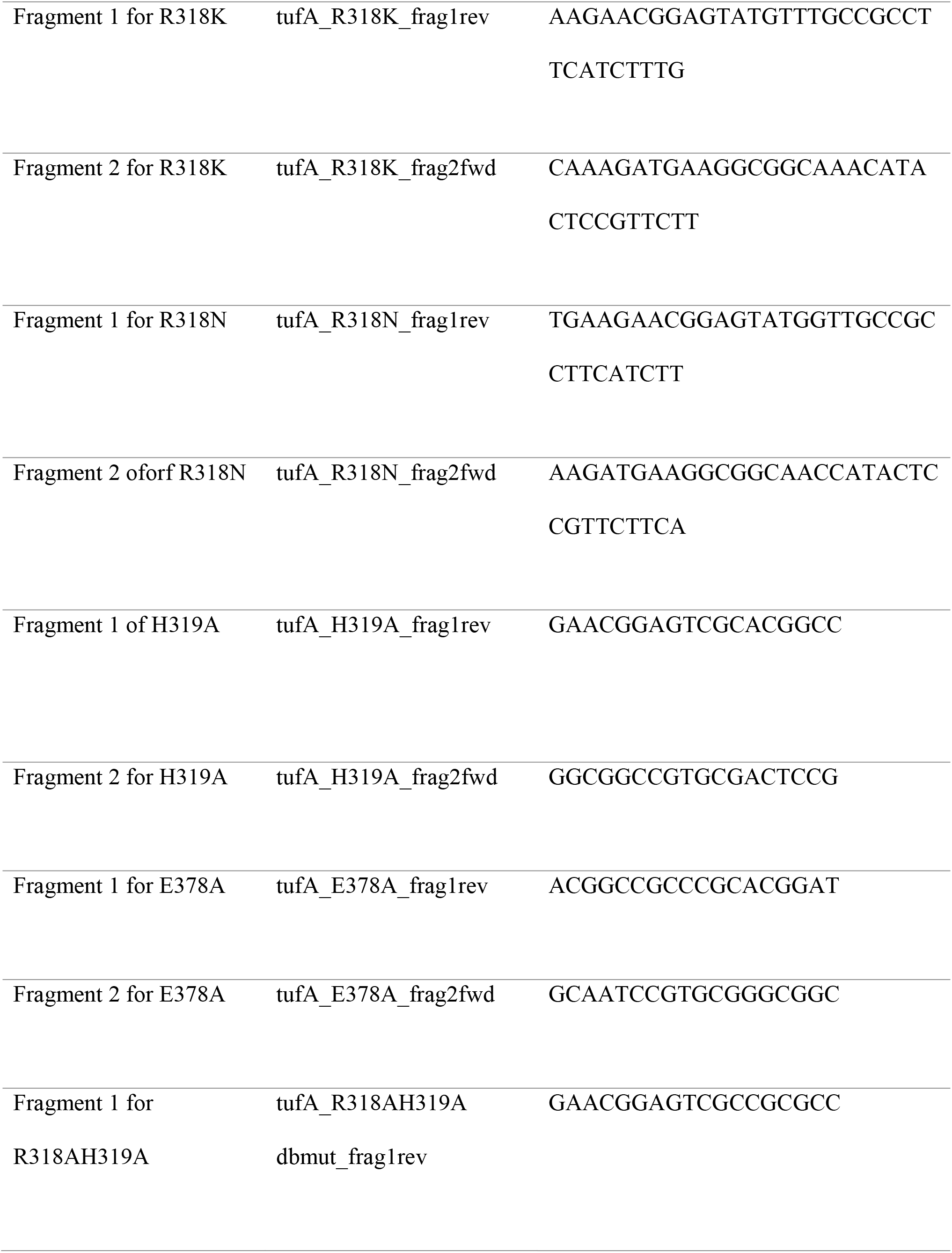
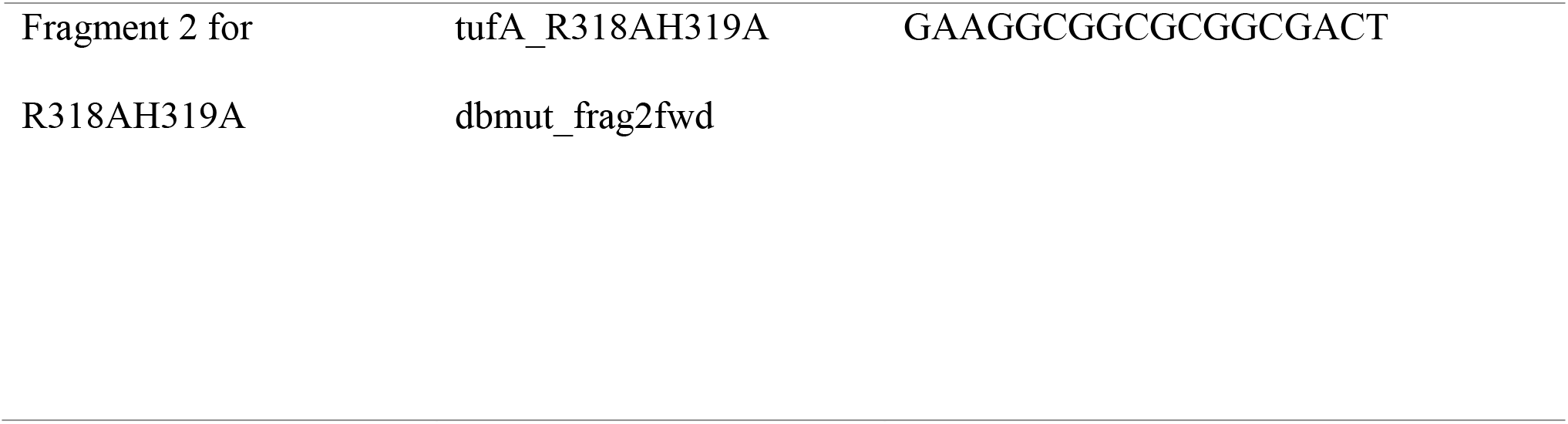
Oligonucleotides used in this study

